# Relational homeostatic scaling supports stable rate-code transmission under noise and heterogeneity

**DOI:** 10.1101/2025.11.25.689806

**Authors:** Huihong Li, Robert A. McDougal

## Abstract

Reliable transmission of firing-rate signals through neural circuits requires synaptic coupling to remain within a narrow regime shaped by neuronal heterogeneity and noise. Outside this regime, classical theory predicts that activity will dissipate or amplify into synchrony. Studies have shown that stable rate-code transmission is biologically achievable, but they leave open how the required operating regime is established. Canonical plasticity rules including Hebbian plasticity and conventional homeostatic scaling do not by themselves recover this regime. We therefore introduce relational homeostatic scaling, a local synaptic scaling rule in which each postsynaptic neuron adjusts its total excitatory afferent strength to reduce the discrepancy between a recent AMPA-weighted afferent activity trace and a recent postsynaptic activity trace. Together, these traces capture the recent rate entering and leaving the neuron without requiring a global rate target. Mean-field analysis and simulations show that this rule drives synaptic weights toward the critical regime without precise weight initialization. The rule remains robust to neuronal heterogeneity, composes with spike-timing-dependent plasticity without disrupting Hebbian competition, and stabilizes the eligibility trace required by three-factor learning rules. These findings suggest that relational homeostatic scaling may provide a substrate for stable rate-code transmission in neural circuits.

**Author Summary:** Neural signals often pass through many processing stages, but this is difficult in networks made of noisy and heterogeneous neurons. Classical feedforward models predict that activity will either fade away or become too synchronized unless synaptic strengths fall within a narrow stable regime. Recent studies have shown that biological neural networks can transmit firing-rate information reliably. How do neural circuits find and maintain the operating regime needed for stable rate-code transmission? We propose a simple local mechanism, relational homeostatic scaling, in which each neuron adjusts the strength of its incoming excitatory synapses by comparing a recent trace of its input activity with a trace of its own activity. Unlike conventional homeostatic scaling, the goal is not to force every neuron toward a fixed firing rate. The goal is to preserve the relationship between input drive and output activity. We show that this rule allows layered spiking networks to self-organize into the stable transmission regime, remains robust to neuronal heterogeneity, and works alongside Hebbian spike-timing plasticity. These results suggest that homeostatic plasticity may help establish the circuit conditions under which experimentally observed rate-code transmission can occur.

## Introduction

Neural systems transmit spiking activity across successive processing stages despite substantial noise and cellular heterogeneity (1–4). Individual neurons differ in membrane time constants, firing thresholds, ion-channel composition, and synaptic release statistics (2, 5). Yet neural circuits maintain signal transfer across multiple regions, from sensory brainstem nuclei through thalamocortical projections to cortical laminae and motor output (6–8). Feedforward networks have provided the canonical abstraction for this process and identified two main transmission modes (1). In one, the signal is carried by synchronous spike packets that maintain submillisecond temporal precision across layers (9–11). In the other, it is carried by graded modulations of firing rate that remain largely asynchronous at the single-neuron level (12–14).

Previous studies on both rate-code and synchrony-code transmission have consistently demonstrated that stable spiking activity capable of preserving stimulus information arises only under specific conditions(1, 15). If effective coupling is too weak, spiking signals decay within a few stages. If it is too strong, correlations grow, activity synchronizes, and the response saturates. Furthermore, the conditions for stable propagation are narrow and sensitive to multiple factors, including connection probability and the pairwise correlations it introduces (1, 16, 17), stochastic neurotransmitter release (18, 19), background noise (20), and short-term synaptic plasticity (21). Which mode dominates depends on the parameter regime and neural systems appear to employ both (1). Biological heterogeneity amplifies this difficulty, because neurons differ in intrinsic and synaptic properties and no single global weight can place every cell at unity gain (2, 3, 5). The required fine-tuning is therefore neuron-specific and cannot rely on the global normalization schemes available to deep artificial networks (22, 23). The gain-control problem is therefore local and depends on information available at each synapse or neuron.

Recent experiments in multilayer cortical cultures challenge idealized feedforward models that predict propagation either dissipates or collapses into synchrony (24). In dense biological networks, synchronized pulse packets evoked a brief transient response that faded across layers and a sustained component that propagated reliably as a rate code through layers (24). In vivo sensory pathways have demonstrated that temporal structure can be converted into rate-coded responses (25, 26). In vivo cortical recordings also show that transient stimuli can evoke tonic or persistent firing in sensory and working-memory circuits (27–29). These findings show that stable rate-code transmission is biologically achievable, but they leave unresolved how synaptic strengths are locally adjusted so that each stage transmits the appropriate drive. Existing evidence suggests that neither excitation–inhibition balance alone nor canonical plasticity rules fully provide a sufficient mechanism for stable propagation (13, 30). Hebbian plasticity can amplify feedforward activity and drive responses toward saturation (30, 31). Classical homeostatic plasticity stabilizes activity around a target firing rate, which can weaken the firing-rate modulations that encode input differences (32–34) . A plasticity mechanism supporting rate-code transmission therefore faces the task of finding the critical gain without erasing rate-code stimulus information or disrupting Hebbian competition.

We introduce relational homeostatic scaling as a local plasticity candidate for establishing the operating regime required for rate-code transmission. Each postsynaptic neuron compares a recent AMPA-weighted afferent activity trace with a recent postsynaptic activity trace and rescales total excitatory afferent strength to reduce their mismatch. Together, these traces capture the recent rate entering and leaving the neuron without defining a separate global input or output rate variable. This rule is motivated by evidence that homeostatic scaling can sense glutamatergic drive (24, 35–38). Meanfield analysis shows that this local relation defines a critical gain regime. Simulations show that the rule moves synaptic weights toward this regime from different initial conditions and compensates for neuronal heterogeneity. The same mechanism preserves rate-code transmission, remains compatible with STDP-based synaptic competition, and stabilizes eligibility traces required by three-factor learning rules. These results identify relational homeostatic scaling as a candidate plasticity mechanism for maintaining stable rate-code transmission in spiking circuits.

## Results

### Feedforward architectures require a finely tuned per-synapse weight for stable rate-code transmission

We first quantified the tuning width of a purely feedforward leaky integrate-and-fire (LIF) circuit that sustains rate-code transmission across depth. We defined stable rate-code transmission as mean-rate reproduction across layers under asynchronous stimulus. In this criterion, population spiking rate is relayed rather than millisecond spike-timing alignment (Fig. 1a).

**Fig 1.**
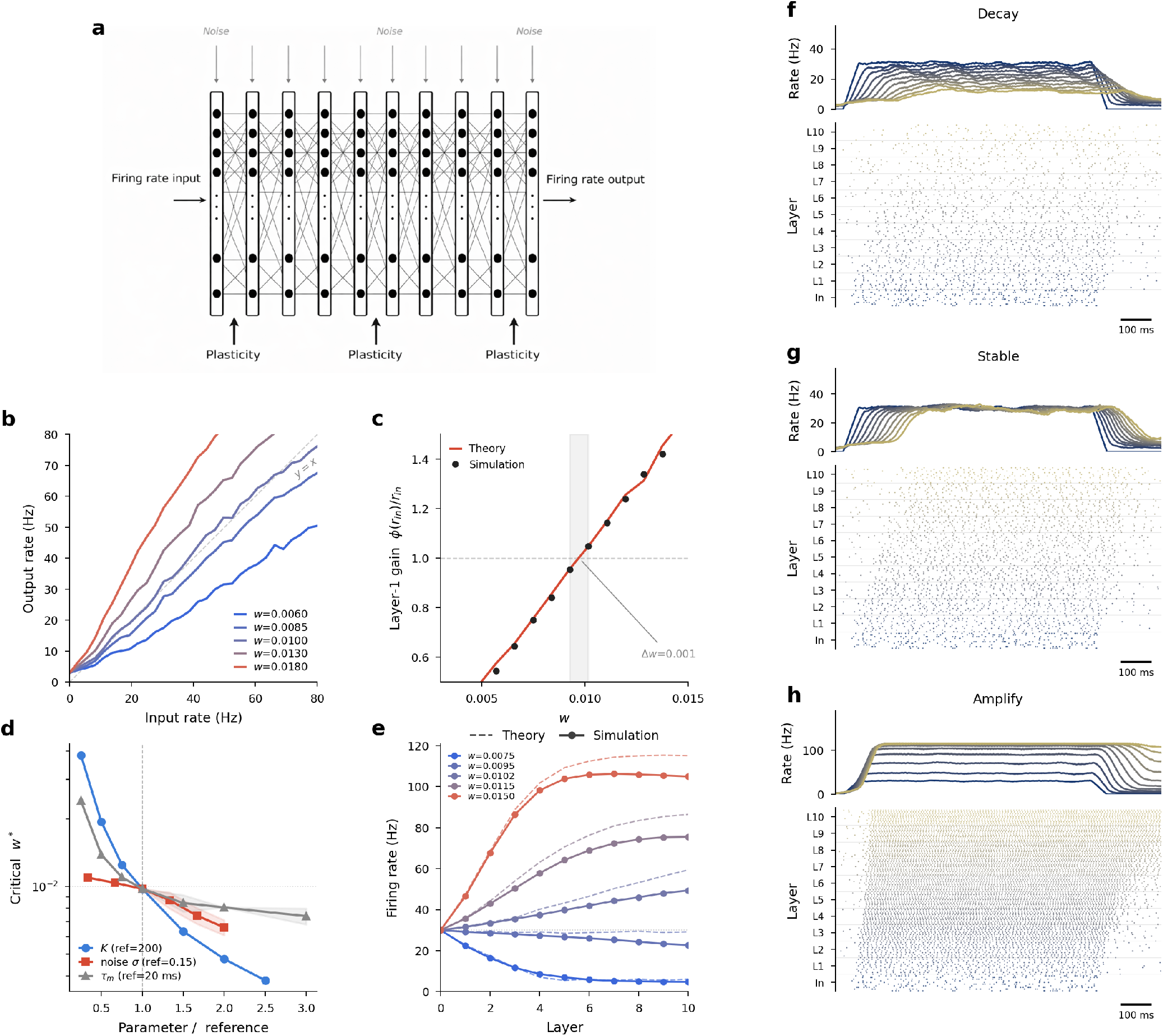
Stable rate-code transmission across a feedforward LIF network requires a finely tuned per-synapse weight. (a) Schematic of the 10-layer feedforward LIF network. Layer In is a 1000-neuron Poisson input layer. Each subsequent layer has 1000 LIF neurons with Bernoulli connectivity (*p* = 0.2, *K* = 200). Every neuron receives independent white-noise current (std. *σ* = 0.15). Synapses in every layer are subject to the plasticity rules described in subsequent sections. (b) Single-neuron LIF transfer function *ϕ*(*r*_in_) obtained by mean-field simulation for a set of raw per-synapse weights (*w* = 0.006–0.018). The grey dashed line is the identity. Only an intermediate *w* places *ϕ* near unit slope at *r*_in_ = 30 Hz. (c) Layer-1 gain *ϕ*(*r*_in_)/*r*_in_ from full network simulation (black circles) vs. mean-field theory (red line). The shaded band highlights the narrow window of *w* in which the gain lies within ± 10% of unity. (d) Critical *w*^∗^ on a shared log axis as a function of fan-in *K*, noise amplitude *σ*, and membrane time constant *τ*_*m*_, each normalized to its reference value (*K* = 200, *σ* = 0.15, *τ*_*m*_ = 20 ms). Shaded envelopes indicate the width of the |gain−1| < 0.1 stability band. (e) Layer-wise mean firing rate across a 10-layer network for five values of *w* bracketing *w*^∗^. The band that keeps *r*_*L*=10_ within ±50% of *r*_in_ extends only from about −5% to +2.5% of *w*^∗^, with the amplify side saturating faster than the decay side cools. (f–h) Per-layer mean firing-rate traces (top, darker = deeper layer) and spike rasters (bottom, 10 randomly sampled neurons per layer) during a 500-ms, 30-Hz Poisson stimulus for three closely spaced choices of *w*: (f) decay (*w* = 0.009), (g) stable (*w* ≈ *w*^∗^ = 0.0095), and (h) amplify (*w* = 0.015).

In a mean-field approximation, each neuron in layer *ℓ* receives a synaptic current whose steady-state mean is 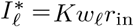, where *K* is the expected fan-in, *w*_*ℓ*_ is the per-synapse weight, and *r*_in_ is the mean firing rate of the preceding layer. The output population rate of the layer is 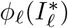, where *ϕ*_*ℓ*_ is the single-neuron transfer function (*f* –*I* curve) set by *τ*_*m*_, *σ*, and the refractory period. Stable rate-code transmission requires two conditions to hold simultaneously. The first is the fixed-point condition

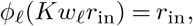

which ensures the mean rate is reproduced at each layer. The second is unit gain at that fixed point, so that rate perturbations are neither amplified nor suppressed. For a small perturbation *δr*_*ℓ*−1_ to the input, the output perturbation is *δr*_*ℓ*_ ≈ *G*_*ℓ*_ *δr*_*ℓ*−1_, where the effective layer gain is

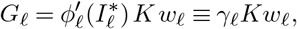

And 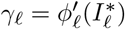 is the slope of the transfer function at the operating point. Chaining *L* layers gives *δr_L_*/*δr*_0_ Π_*ℓ Gℓ*_, so a fractional deviation *ε* in *G*_*ℓ*_ amplifies exponentially with depth as 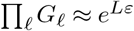. Setting *G*_*ℓ*_ = 1 at the fixed-point operating current defines a critical per-synapse weight

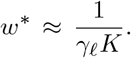

Because *γ*_*ℓ*_ is the *f*–*I* slope evaluated at the operating point, *w*^∗^ is set jointly by fan-in *K*, noise amplitude *σ*, and membrane time constant *τ*_*m*_ (all defined in the simulation paragraph below), and no universal setting exists.

To test this prediction we simulated 10-layer feedforward stacks of *N* = 1000 LIF neurons per layer, driven by a 1000-neuron Poisson input firing at *r*_in_ = 30 Hz. Each postsynaptic cell received sparse Bernoulli afferents with connection probability *p* = 0.2 and fan-in *K* = *N*_in_ *p* = 200, parameterised by per-synapse weight *w*, so that the total excitatory drive onto a cell scaled as *wKr*_in_ per ms. Intrinsic fluctuations were modelled as additive white noise of amplitude *σ* = 0.15 on a *τ*_*m*_ = 20 ms membrane (Fig. 1a). The single-neuron transfer function *ϕ*(*r*_in_) interpolated between subthreshold compression and near-linear gain across *w* (Fig. 1b), and layer-1 simulation rates tracked the mean-field theory, crossing unit gain at *w*^∗^ ≈ 0.0098 (Fig. 1c). Sweeping *K, σ*, and *τ*_*m*_ over a factor of three moved *w*^∗^ by nearly an order of magnitude, and the band of weights for which layer gain stayed within ±10% of unity was narrow in every case (Fig. 1d). *w*^∗^ fell as 1/*K* and also decreased with both *σ* and *τ*_*m*_, consistent with *w*^∗^ ∝ 1/(*γK*).

Even with all other parameters fixed, the precision required was stringent and asymmetric. Sweeping *w* in 2.5% steps and iterating mean-field rates through ten layers (Fig. 1e), a +5% deviation drove the layer-10 rate above 60 Hz, and a −10% deviation collapsed it below 10 Hz. Keeping *r*_*L*=10_ within ±50% of *r*_in_ required *w* to lie within roughly [−5%, +2.5%] of *w*^∗^. The amplify side was the more sensitive, because the LIF *f*–*I* curve is supralinear above threshold and a small excess drive recruits extra spiking that compounds across depth. On the decay side the layer-10 rate falls more gradually and then settles onto a ∼3–4 Hz floor set by spontaneous threshold crossings. Three representative operating points (Fig. 1f–h) show the corresponding spiking dynamics. At *w* = 0.009 population activity fails to reach the deep layers (*decay*), at *w* ≈ *w*^∗^ the 30-Hz signal is relayed across all ten layers (*stable*), and at *w* = 0.015 firing saturates after a few layers (*amplify*). Because *w*^∗^ depends on cellular and network parameters that are themselves variable, this condition is unlikely to be met or maintained without an active compensatory mechanism.

### Canonical plasticity rules fail to recover the stable transmission regime

Local plasticity may establish the operating regime needed for rate-code transmission without a globally tuned weight. Hebbian plasticity and homeostatic synaptic scaling are the main local candidates for this task. We next asked whether canonical variants of these rules can recover the narrow critical band identified in Fig. 1. We crossed three initial weight regimes (*w*_0_ = 0.0050, 0.0095, 0.0150) with five STDP variants and five homeostatic conditions. The criterion was the mean layer-wise firing-rate error 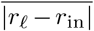 over 100 training epochs (Fig. 2, see Methods).

**Fig 2.**
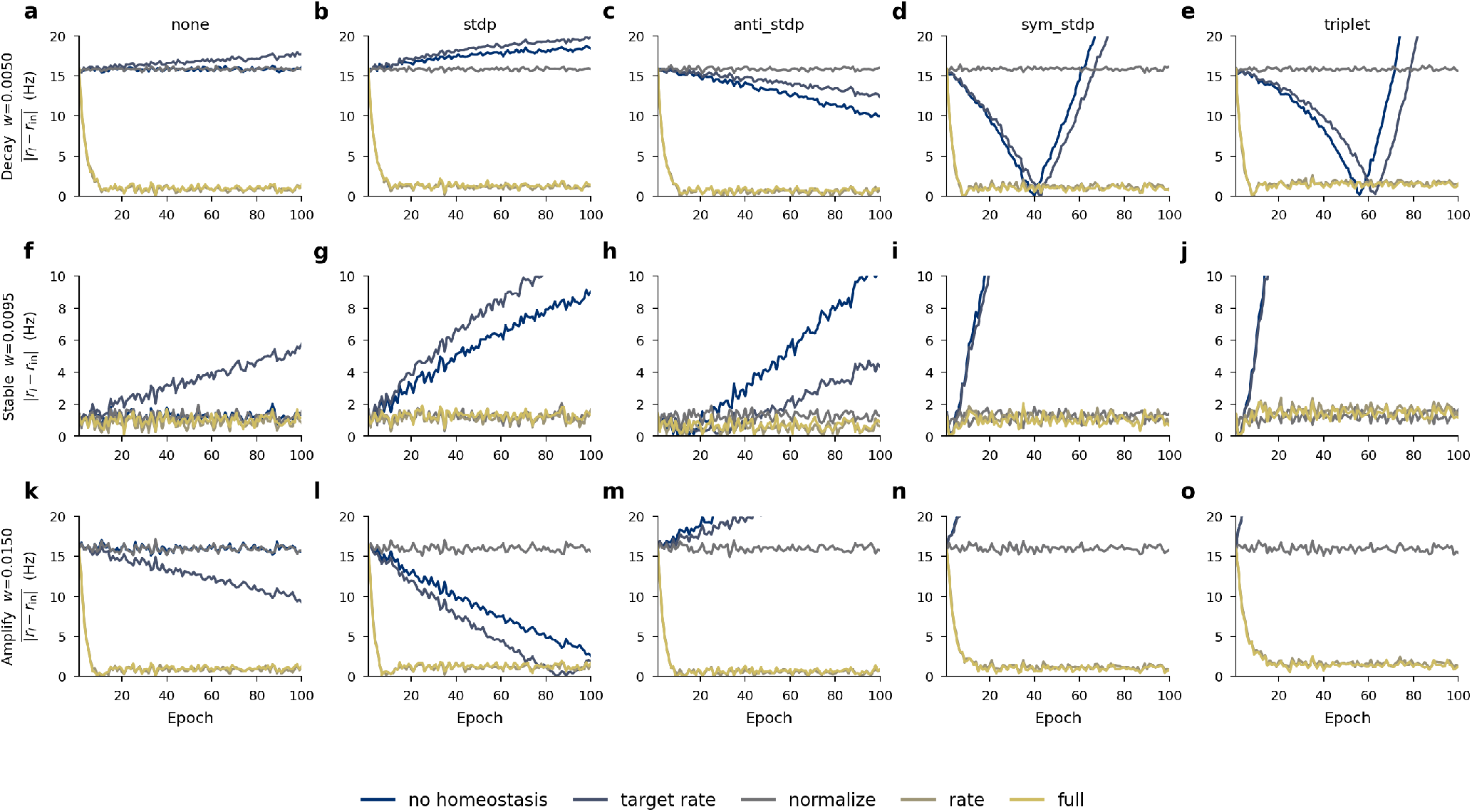
Neither Hebbian nor classical homeostatic rules recover stable rate-code transmission. The first layer of the feedforward stack (*K* = 200) was initialized in each of the three weight regimes identified in Fig. 1 (rows, *decay w*_0_ = 0.0050, *stable w*_0_ = 0.0095, *amplify w*_0_ = 0.0150). For every initialization we crossed five STDP variants (columns, from no STDP through triplet STDP) with five homeostatic rules (line colors, no homeostasis, target-rate fixpoint, total-weight normalization, the proposed relational rule 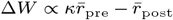, and the combined rule). Each curve is the mean absolute layer-wise firing-rate error 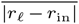 over 100 training epochs (500 ms stimulus at 30 Hz per epoch). Without homeostasis, every STDP variant increases the error from every initial regime, most severely for symmetric STDP. Target-rate homeostasis attenuates the signal even when the network begins in the stable regime. Total-weight normalization holds the stable regime stable but locks the decay and amplify regimes in place. The relational rule and its combined variant drive the error to zero from every initialization.

Acting without homeostasis, every STDP variant pushed synaptic weights upward, driving the network toward the amplify regime from every initialization (grey curves, Fig. 2). The mechanism is the causal timing bias of feedforward architectures. Presynaptic spikes systematically precede post-synaptic spikes, so any STDP kernel that is net-potentiating on causal coincidences accumulates weight monotonically. From the decay and stable initializations this caused rates to overshoot *r*_in_ and the error to grow in every variant. Symmetric STDP was the most destabilizing because it potentiates on both causal and acausal coincidences, so the total afferent weight grew unconditionally and the error diverged from every starting point (Fig. 2d,i,n).

Classical homeostatic rules failed for complementary reasons. A fixed-target rule drove each neuron toward a preset rate rather than toward the input rate, so any stimulus-driven rate deviation was actively attenuated back to the pre-set level, preventing rate-code signals from being relayed (Fig. 2f, gold curve). This failure persisted even from the stable initialization, because the preset target did not in general coincide with *r*_in_. Total-weight normalization preserved the initial operating point, keeping the stable regime stable (Fig. 2f) but freezing the decay and amplify regimes in their failure modes (Fig. 2a,k). Normalization is conservative and cannot move an ill-initialized circuit into the stable band.

Every STDP variant destabilized rate-code transmission when acting alone, and neither classical homeostatic rule could both discover and maintain the stable band across initializations. We therefore introduce a local relational rule whose fixed point aligns recent AMPA-weighted afferent activity with recent postsynaptic activity rather than a preset target rate. In the same sweep, the relational rule and its combination with a target-rate term drove the error to zero from every initialization and for every STDP variant (Fig. 2). These failures reflect a mismatch between the controlled variable and the computational requirement of rate-code transmission. STDP controls relative spike timing, fixed-target scaling controls absolute firing rate, and total-weight normalization fixes total afferent weight at its initial value. Stable rate-code transmission requires control of the relationship between afferent activity and postsynaptic activity. We analyze this rule in the remaining sections.

### Relational homeostatic scaling recovers the critical weight across parameters

Stable rate-code transmission requires homeostasis to compare the recent afferent activity trace with the recent postsynaptic activity trace. A fixed postsynaptic firing-rate target can stabilize activity, but it does not encode whether the current output preserves the incoming rate signal. We therefore asked whether a relational version of synaptic scaling can use the AMPA-weighted afferent activity trace and a postsynaptic activity trace to discover the critical weight. The rule augments classical scaling with this AMPA-weighted afferent activity trace. Experiments across multiple neural circuit types support the biological plausibility of this added afferent component (24, 35–38). The total excitatory weight *W*_*j*_ onto postsynaptic neuron *j* evolves as

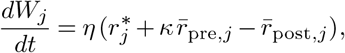

where 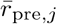 is the AMPA-weighted afferent activity trace, computed as a weight-normalized low-pass filter of the presynaptic spike train. 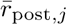 is a low-pass filter of the neuron’s own firing, and 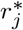 is its intrinsic homeostatic target. Setting *κ* = 0 recovers the classical synaptic scaling rule. In the simulations below, we assume 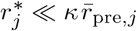 at the firing rates studied, so the intrinsic target contributes negligibly and the rule reduces to 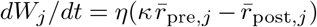. The update is implemented as a multiplicative rescaling of all excitatory afferents onto neuron *j*, so that *W*_*j*_ → *W*_*j*_ +Δ*W*_*j*_ while preserving the relative strength of individual synapses (Fig. 3a). The fixed point aligns the neuron’s recent AMPA-weighted afferent activity trace with its recent postsynaptic activity trace, scaled by *κ*. This allows the rule to capture the recent rate entering and leaving each neuron without imposing a fixed global rate target. Linearizing around this fixed point shows that deviations relax as 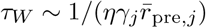. Learning is faster at higher presynaptic activity or single-neuron gain, and averaging over long enough traces makes the rule insensitive to millisecond-scale fluctuations.

**Fig 3.**
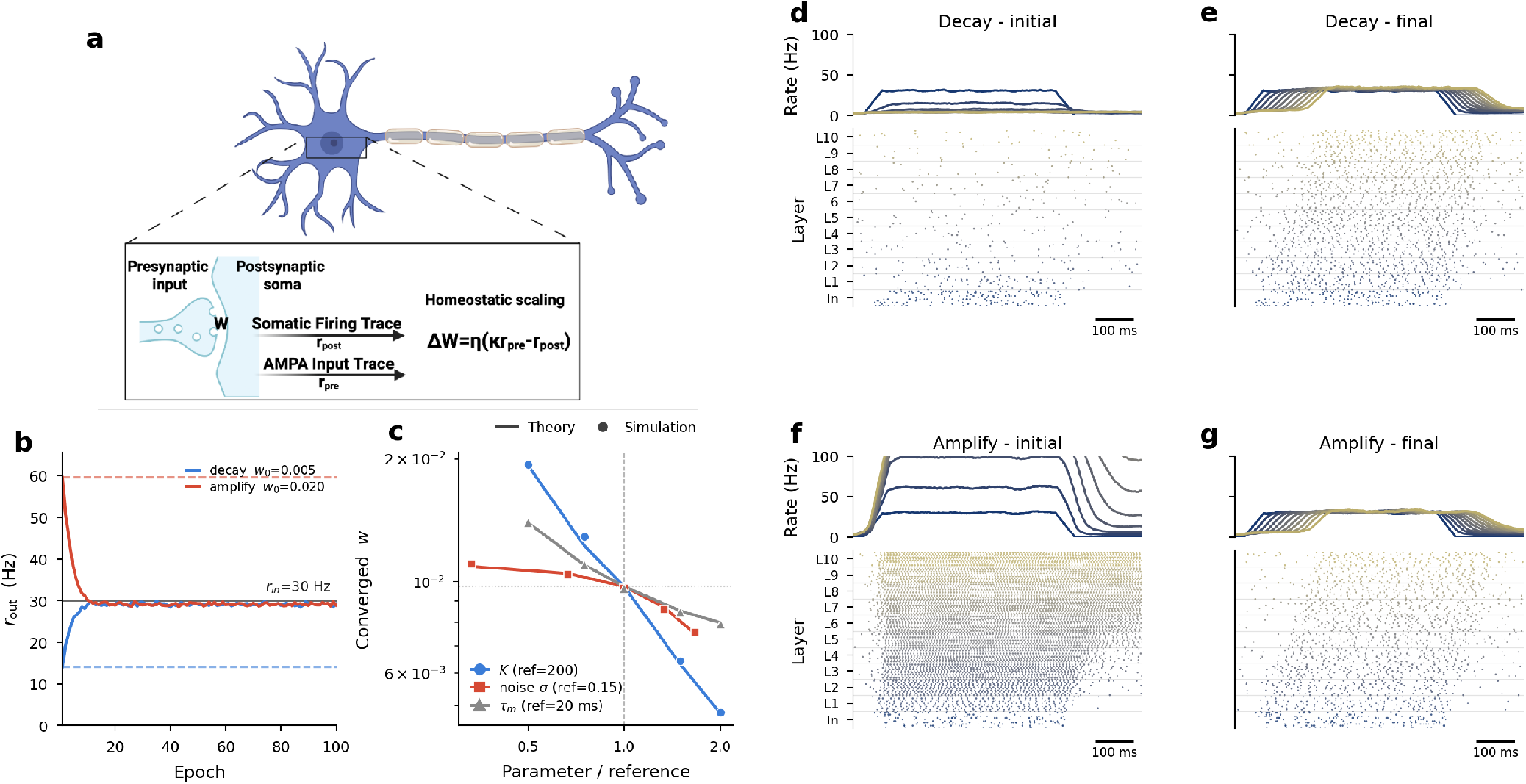
Relational homeostatic scaling drives networks back to the critical weight *w*^∗^. (a) Schematic of the local learning rule. Each postsynaptic neuron maintains an AMPA-weighted afferent activity trace 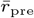 and a postsynaptic activity trace 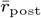. Its total excitatory weight *W* is updated by 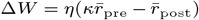, so the fixed point aligns the two recent activity traces after scaling by *κ*. (b) Single-layer mean output rate over 100 training epochs starting from *decay* (*w*_0_ = 0.005, blue) and *amplify* (*w*_0_ = 0.020, red) initial weights. Both trajectories converge to *r*_in_ = 30 Hz (solid black line), while dashed horizontals mark the pre-training rates. (c) Converged per-synapse weight *w* after training matches the theoretically predicted *w*^∗^ (solid lines) across sweeps of fan-in *K* (blue), noise *σ* (red), and *τ*_*m*_(grey), each normalized to its reference. Markers show simulation-derived converged weights (circles *K*, squares *σ*, triangles *τ*_*m*_). (d–e) Layer-wise rate trace (top) and raster (bottom) of a 10-layer network initialized in the decay regime, shown *initial* (pre-training, d) and *final* (after 100 epochs, e). The initial network fails to relay activity past the input layer, while the trained network achieves near-unity inter-layer gain through all ten layers (per-stage attenuation below 4%). (f–g) Same for the amplify initialization (*w*_0_ = 0.020), where the initial network saturates and the trained network recovers stable layer-wise rate-code transmission with the same per-stage gain accuracy.

A single LIF layer initialized in either the decay (*w*_0_ = 0.005) or the amplify (*w*_0_ = 0.020) regime of Fig. 1 reliably converged to the target rate within ∼30 epochs (Fig. 3b). The converged per-synapse weight tracked the theoretically predicted *w*^∗^ across three independent sweeps, scaling inversely with fan-in *K* and decreasing with both noise amplitude *σ* and membrane time constant *τ*_*m*_ (Fig. 3c). In every sweep the simulation markers fell on the mean-field theory curves, confirming that the rule discovers the same *w*^∗^ identified by the rate analysis.

The 30-Hz training protocol does not by itself show whether the recovered weight is specific to the training rate or whether it places the layer in the linear operating regime identified in Fig. 1b,c. We therefore froze a single layer after 100 epochs of relational scaling from the decay initialization and changed only the Poisson input rate from 5 to 80 Hz. The trained layer reproduced the input rate across the full sweep, with frozen-test gain ranging from 0.90 to 1.11 and gain 0.98 at 30 Hz (Supplementary Fig. S1). This confirms that the learned weight lies on the near-linear transfer-function segment found in Fig. 1, rather than a rate-specific solution for the training stimulus. Training at 30 Hz therefore identifies an operating regime suitable for transmission of other rate-coded inputs within the tested range.

Extending the analysis to a 10-layer network, we trained the decay and amplify regimes for 100 epochs. The initial decay network failed to relay activity past layer 1 (Fig. 3d). After training, all ten layers transmitted a near-target signal, with the inter-layer gain deviating by less than 4% from unity at each stage (Fig. 3e). Symmetrically, the initial amplify network saturated at every layer (Fig. 3f), and training restored layer-wise rate-code transmission with comparable inter-layer gain accuracy across depth (Fig. 3g). The same local rule therefore simultaneously suppressed runaway amplification and recovered stable rate-code transmission from the decay regime, using only information available at each postsynaptic neuron.

### Relational homeostatic scaling is robust to neuronal heterogeneity

Biological heterogeneity makes the critical operating point neuron specific. Neurons differ in intrinsic excitability and membrane time constant across species, brain regions, and cell types. We therefore asked whether relational homeostatic scaling can preserve population-level rate-code transmission under broad cell-to-cell variability. We modeled heterogeneity either as variation in the gain factor *κ*_*j*_ or as variation in *τ*_*m,j*_. The relational fixed point is 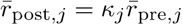. The population mean therefore remains aligned with the input rate when ⟨*κ*_*j*_⟩ = 1, while each neuron adopts its own total afferent weight (Fig. 4).

**Fig 4.**
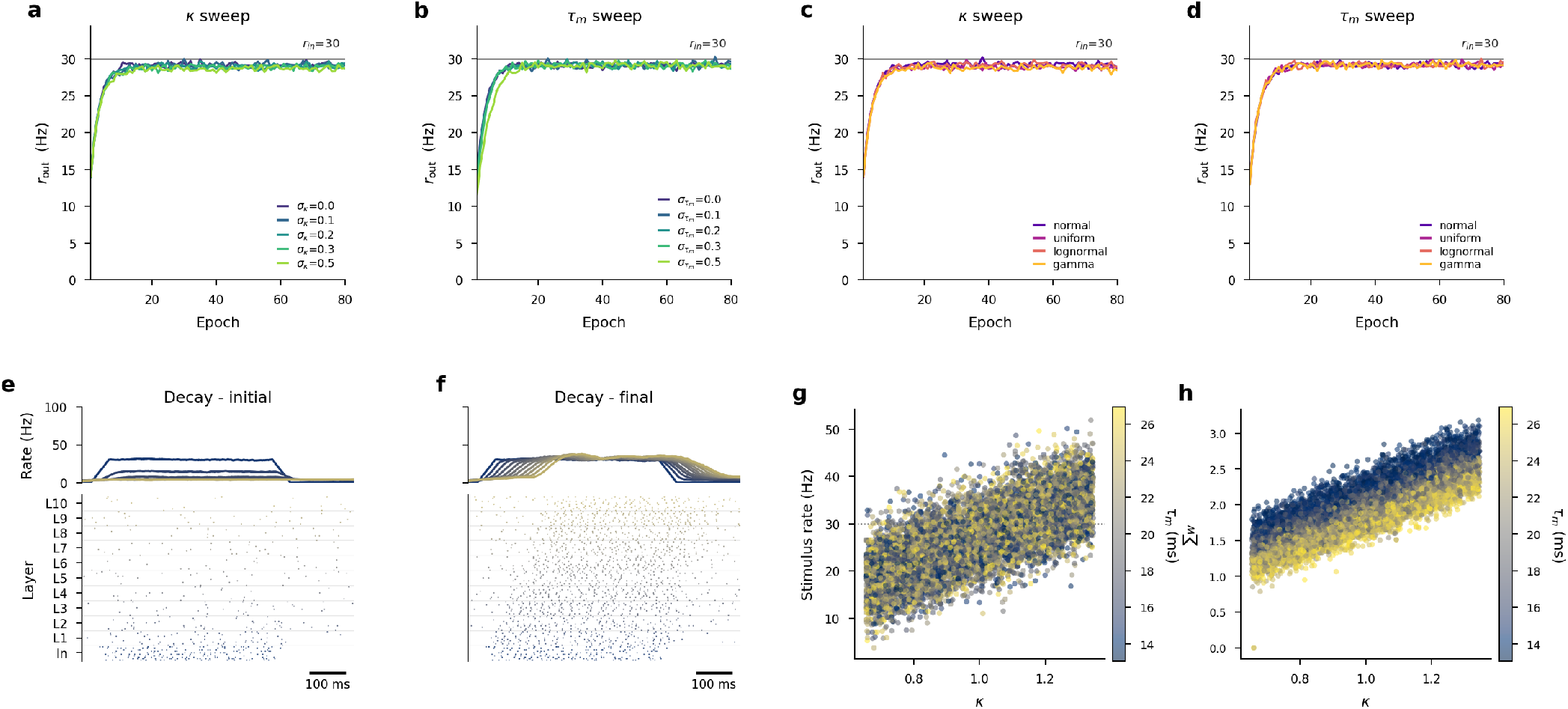
Relational homeostatic scaling tolerates broad neuronal heterogeneity. (a–b) Single-layer mean output rate *r*_out_ over 80 training epochs when each neuron’s (a) gain factor *κ* or (b) membrane time constant *τ*_*m*_ is drawn from a uniform distribution with coefficient of variation *σ* ∈{0, 0.1, 0.2, 0.3, 0.5} (lighter color = larger spread). The population mean converges to *r*_in_ = 30 Hz for every spread. (c–d) Same analysis at a fixed spread *σ* = 0.3, with heterogeneity drawn from four different distribution families (normal, uniform, lognormal, gamma), applied independently to (c) *κ* and (d) *τ*_*m*_. The rule converges under every distribution family. (e–f) 10-layer decay network (*w*_0_ = 0.005) with moderate heterogeneity ( 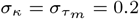, uniform), shown *initial* (e) and *final* (f). Training restores rate-code transmission through all layers despite per-neuron variability. (g) Per-neuron stimulus firing rate in the final network as a function of *κ*, colored by *τ*_*m*_. Firing rates align with the *κr*_in_ line predicted by the rule, independent of *τ*_*m*_. (h) Total afferent weight *W*_*j*_ per neuron vs. *κ*, same color code. Each neuron adopts a different *W*_*j*_ that exactly compensates for its individual gain, in line with 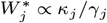.

To test this, we trained single-layer networks in the decay regime, sweeping the coefficient of variation of either *κ*_*j*_ or *τ*_*m,j*_ across *σ* ∈ {0, 0.1, 0.2, 0.3, 0.5} under a uniform distribution. For every spread, the population-mean output rate converged to *r*_in_ = 30 Hz within 80 training epochs (Fig. 4a– b), including at *σ* = 0.5, where individual neurons span a two-fold range in gain. At a fixed spread of *σ* = 0.3, we varied the distribution family across normal, uniform, lognormal, and gamma. All four families produced indistinguishable convergence (Fig. 4c–d), suggesting that the relevant quantity is the population mean of *κ*_*j*_, not the parametric form of its distribution.

We then extended to a 10-layer network with moderate heterogeneity in both *κ*_*j*_ and *τ*_*m,j*_ (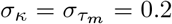, uniform, ⟨*κ*_*j*_⟩ = 1) starting from the decay regime. The untrained network failed to relay activity beyond the input layer (Fig. 4e). After 100 training epochs, every layer transmitted a near-target signal with per-stage attenuation below 3%, comparable to the homogeneous case (Fig. 4f). Each neuron’s stimulus-window firing rate aligned with the prediction *r*_*j*_ = *κ*_*j*_*r*_in_ with negligible dependence on *τ*_*m,j*_ (Fig. 4g), so the population mean satisfied ⟨*κ*_*j*_⟩ *r*_in_ = *r*_in_ by construction. The total afferent weight *W*_*j*_ rose with *κ*_*j*_ and scattered with *τ*_*m,j*_ (Fig. 4h), consistent with 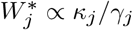. Each neuron autonomously compensated for its own intrinsic properties while the population preserved stable rate-code transmission.

### Relational homeostatic scaling is compatible with spike-timing-dependent plasticity

A useful circuit preserves rate-code transmission while synapse-specific learning selects which afferents dominate the transmitted signal. Hebbian STDP can create broad, heavy-tailed afferent weight distributions under homeostatic constraints (33, 34). We next asked whether relational homeostatic scaling preserves this synapse-level competition. We treated total afferent weight and relative synaptic weights as separate degrees of freedom, and evaluated five STDP variants with the relational rule active (Fig. 5).

**Fig 5.**
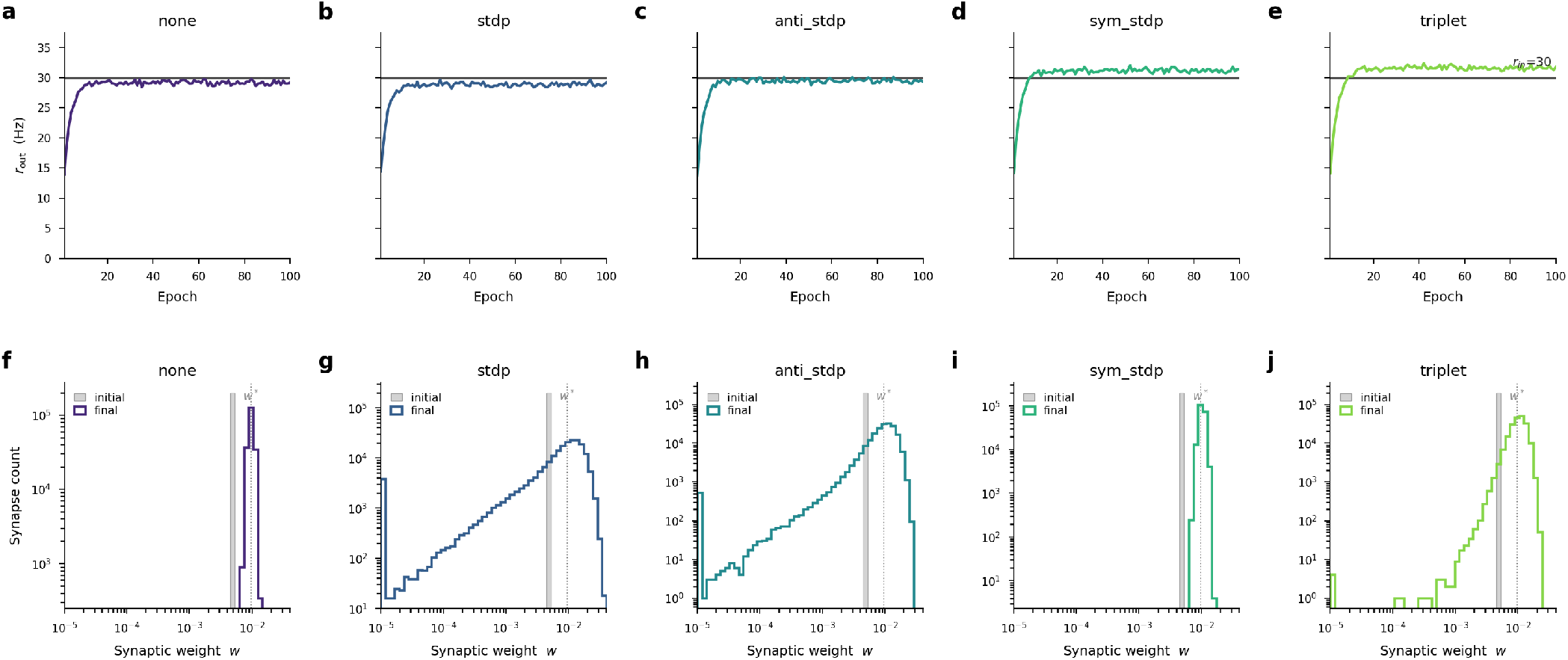
Relational homeostatic scaling composes cleanly with the tested STDP variants. The feedforward network was trained from the decay initialization (*w*_0_ = 0.005) with the relational rule active. Afferent weight organization at the first layer is shown for each of the five STDP variants. (a–e) Mean output rate *r*_out_ at the first layer over 100 epochs for (a) no STDP, (b) pair-based STDP, (c) anti-STDP, (d) symmetric STDP, and (e) triplet STDP. All variants converge to *r*_in_ = 30 Hz (solid black). (f–j) Corresponding log-log histograms of per-synapse afferent weights at the first layer before (filled grey) and after (colored step) training. Homeostasis-only training (f) shifts the delta distribution to *w*^∗^ without broadening it, while pair-based, anti-causal, and triplet STDP (g,h,j) broaden the distribution across two orders of magnitude with a fraction of synapses silenced and a heavy tail of potentiated afferents. The mean per-synapse afferent weight (grey vertical line, *w*^∗^) remained at the critical value in every case.

Starting from the decay initialization (*w*_0_ = 0.005), we trained the feedforward network with the relational rule active and recorded the afferent weight distributions that developed at the first layer under no STDP, pair-based, anti-causal, symmetric, and triplet STDP (Fig. 5). The population-mean output rate converged to *r*_in_ = 30 Hz in every condition (Fig. 5a–e), suggesting that the total-weight constraint holds independently of which STDP kernel is active. The same result held across all three weight initializations and all STDP variants in the systematic comparison of Fig. 2, where every other homeostatic rule failed to reach zero error under at least one condition.

The weight distributions (Fig. 5f–j) reveal the division of labor. Homeostasis alone shifted the initial delta distribution from *w*_0_ = 0.005 to *w*^∗^ without broadening it (Fig. 5f), consistent with a pure total-weight rescaling. Adding pair-based or triplet STDP produced a qualitatively different outcome, with weights spanning two orders of magnitude, a fraction of afferents silenced (weight near zero), and a heavy tail of potentiated synapses extending up to four times *w*^∗^ (Fig. 5g,j), consistent with the heavy-tailed distributions reported when STDP operates under homeostatic constraints (33, 34). Anti-causal STDP produced a similar heavy-tailed structure (Fig. 5h). Symmetric STDP, which potentiates on both causal and acausal coincidences, created no directional competitive pressure and left the distribution near its initial shape (Fig. 5i). In every case the mean per-synapse afferent weight remained at *w*^∗^, so the total afferent weight stayed at the value required to relay *r*_in_ while the per-synapse distribution reflected the synapse-specific structure imposed by each STDP variant.

Relational homeostatic scaling therefore constrains the total afferent weight per neuron without interfering with per-synapse Hebbian competition. The two rules target orthogonal degrees of freedom of the weight matrix. Relational homeostatic scaling controls the total afferent weight per neuron while STDP redistributes individual synaptic strengths, so both can operate simultaneously without eroding each other’s functional outcome. The network simultaneously maintains stable layer-to-layer rate-code transmission and preserves the broad, graded distribution of afferent strengths that STDP-driven Hebbian competition produces in biological circuits (33, 34). In this division of labor, relational homeostatic scaling sets the operating point, STDP selects the pathway, and eligibility traces provide the local substrate on which delayed teaching signals can act.

### Relational homeostatic scaling stabilizes eligibility traces for local synaptic credit assignment

Rate-code transmission also sets the local activity statistics used by downstream credit-assignment rules. Three-factor plasticity gates a local eligibility trace with a delayed teaching signal, such as a neuromodulatory broadcast or a top-down error. Deep spiking learning rules such as e-prop (eligibility propagation) and PP-prop (pre-post eligibility trace propagation) require eligibility amplitudes that remain comparable across layers (39, 40) (see Methods). We asked whether relational homeostatic scaling stabilizes this eligibility trace across depth. We measured the eligibility trace amplitude during training from decay, stable, and amplify initializations (Fig. 6).

**Fig 6.**
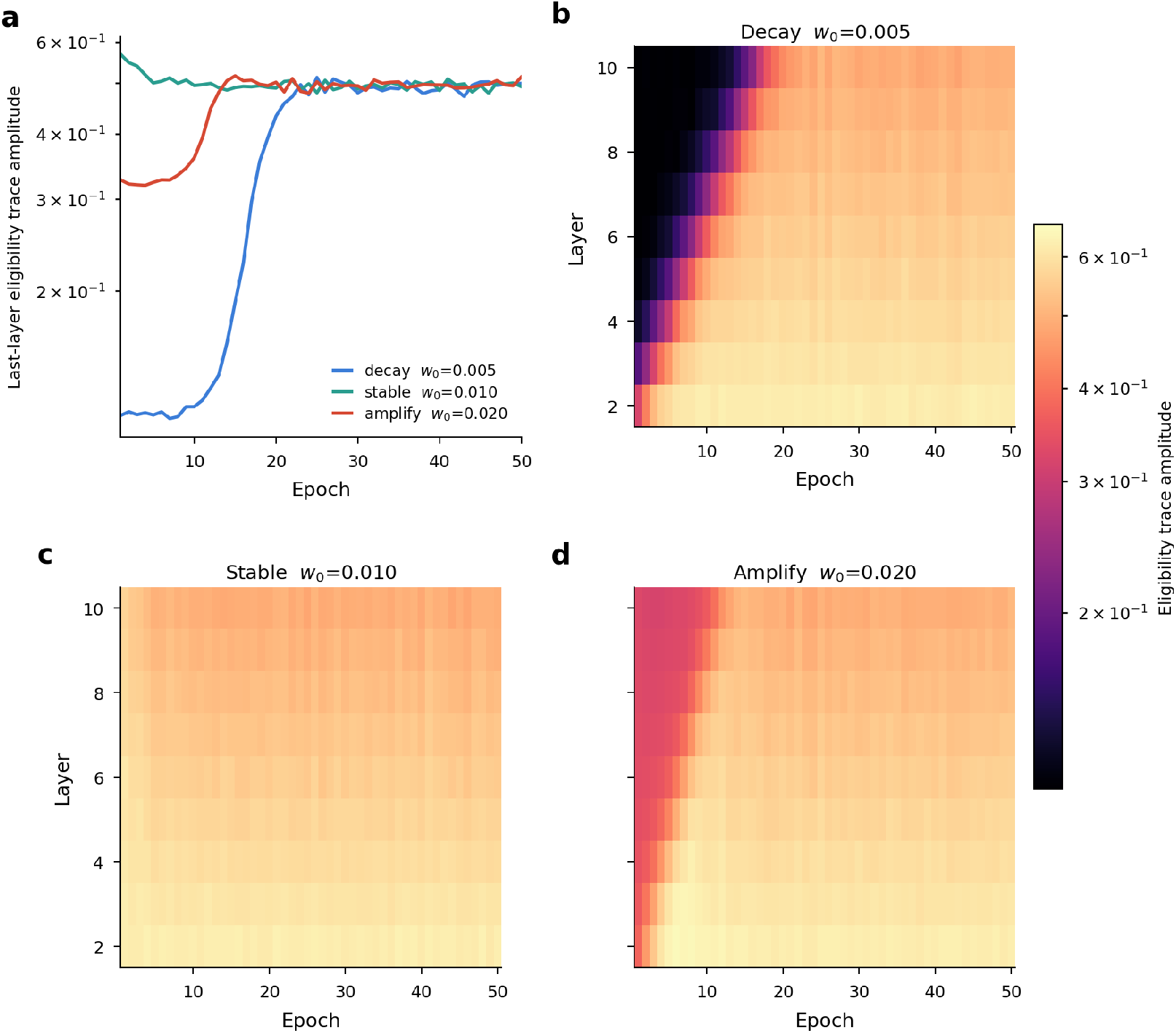
Relational homeostatic scaling equalizes the per-layer eligibility trace that local credit-assignment rules depend on. A 10-layer network was trained with the relational rule from each of the three regimes of Fig. 1 (*decay w*_0_ = 0.005; *stable w*_0_ = 0.010; *amplify w*_0_ = 0.020). At every training epoch the per-layer eligibility-trace amplitude ∥*ϵ*_*f*_ ∥∥*ϵ*_*I*_ ∥ was recorded. This is the quantity that gates a downstream teaching signal in three-factor credit-assignment schemes such as e-prop (eligibility propagation) and PP-prop (pre-post eligibility trace propagation). (a) Last-layer eligibility-trace amplitude over epochs for the three regimes. Decay starts approximately five-fold below the stable trajectory. The amplify regime also starts below the stable level because network-wide saturation suppresses the surrogate-gradient filter, and both converge to it within ∼ 25 epochs. (b–d) Layer-vs-epoch heatmaps of eligibility-trace amplitude (layer 2 at bottom to layer 10 at top) for the decay, stable, and amplify initializations. The decay regime has vanishing trace in deep layers (dark tiles), the amplify regime has uniformly depressed trace across all layers, and the stable regime is spatially uniform. Under the relational rule every initialization converges to the stable profile.

The per-synapse eligibility trace amplitude is ∥*ϵ*_*f*_∥ ∥*ϵ*_*I*_∥, where *ϵ*_*I*_ is a low-pass filter of afferent synaptic current and *ϵ*_*f*_ a low-pass filter of the surrogate activation encoding proximity to threshold. In the decay regime, silent deep layers suppress *ϵ*_*f*_ throughout those layers, so the trace amplitude at layer 10 is approximately five-fold smaller than at layer 2 (Fig. 6b). In the amplify regime, neuronal saturation suppresses the surrogate-gradient filter *ϵ*_*f*_ network-wide, depressing the eligibility-trace amplitude below the stable level throughout the depth of the network (Fig. 6d). Only the stable regime yields a spatially uniform trace at the expected amplitude across all ten layers (Fig. 6c).

With the relational rule active, both pathological initializations converged onto the stable trace profile within ∼25 epochs (Fig. 6a). The per-layer heatmaps confirm that the decay regime’s vanishing deep-layer trace and the amplify regime’s globally depressed trace were corrected, leaving an amplitude profile comparable to the stable initialization (Fig. 6b,d). By anchoring each layer’s mean firing rate to the input rate, the rule simultaneously sets a consistent proximity-to-threshold at every layer, equalizing the eligibility product across depth and providing a depth-stable substrate on which downstream teaching signals can gate comparable local eligibility traces.

Collectively, the results show that a single biologically plausible, per-neuron rule (i) recovers the critical per-synapse weight *w*^∗^ identified by the mean-field analysis, (ii) maintains stable rate-code transmission across feedforward networks regardless of initialization, (iii) tolerates broad per-neuron heterogeneity while preserving the population-level relation between AMPA-weighted afferent activity and postsynaptic activity, (iv) coexists with each of the five tested STDP variants without destroying its heavy-tailed weight signature, and (v) equalizes the per-layer eligibility trace amplitude across depth, providing a stable substrate for local synaptic credit assignment on top of Hebbian competition.

## Discussion

Spiking circuits can transmit rate-code signals through noisy and heterogeneous neural populations. Our results show that this requires each effective synaptic coupling to remain near a narrow critical value set by cellular heterogeneity, membrane noise, and network architecture (Fig. 1). Neither Hebbian nor classical homeostatic rules maintain this precision across initial conditions (Fig. 2). We propose relational homeostatic scaling as a local rule that multiplicatively rescales each neuron’s total afferent weight by the mismatch between a recent AMPA-weighted afferent activity trace and a recent postsynaptic activity trace. The fixed point aligns AMPA-weighted afferent activity with postsynaptic activity rather than enforcing a preset firing-rate target. The rule converges to the theoretically predicted critical weight from any initialization. It also tolerates broad per-neuron heterogeneity, preserves STDP-driven synaptic competition, and equalizes the per-layer eligibility-trace amplitude required for downstream credit assignment (Figs. 3–6). A single local plasticity rule can therefore self-organize feedforward circuits into the operating regime required for stable rate-code transmission.

Prior analyses of feedforward networks have shown that stable rate-code transmission requires synaptic coupling to sit within a narrow critical band, typically enforced by global parameter tuning or E/I-balanced inhibition (1, 9, 13, 14). E/I balance has been the leading proposed mechanism, because inhibitory interneurons absorb excess excitatory drive and extend the stable operating range. Yet even in exactly balanced networks, rate codes deteriorate in long feedforward chains as correlated activity builds and synchronous bursts emerge within a few stages (1, 13). Membrane noise can ameliorate this by linearizing single-neuron transfer functions (20). Per-neuron variability in gain and time constant still shifts the critical weight at each cell individually. Hebbian plasticity then continuously displaces the operating point because the causal timing bias of feedforward connectivity accumulates net potentiation toward saturation (41, 42). Classical homeostatic rules attenuate the relayed signal by targeting a preset rather than a presynaptically defined firing rate (32–34). Our results show that parameter sensitivity, cellular heterogeneity, and Hebbian drift can be resolved together by a single local rule. The failures of existing local rules reflect the fact that these rules regulate variables other than the afferent-postsynaptic relation required for stable rate-code transmission.

The AMPA-weighted afferent activity trace has experimental support, although the molecular pathway is not established. Homeostatic synaptic scaling is mediated by regulated insertion of GluA1-containing AMPA receptors. PKA-AKAP5 scaffolding and GluA1-Ser845 phosphorylation mediate bidirectional scaling (36). Glial TNF-*α* and Arc coordinate synapse-wide scaling (43). Postsynaptic calcium and CaMKII integrate sustained activity changes and translate them into adjustments of AMPA receptor surface density. Classical formulations assume a cell-autonomous firing-rate set point (36, 44). Experiments show that upward synaptic scaling can be triggered by reduced glutamatergic neurotransmission rather than reduced postsynaptic spiking alone (38, 45). This implies that the postsynaptic cell senses AMPA-weighted afferent activity as part of its homeostatic signal (38). A presynaptic-dependent scaling formulation also produces more stable network dynamics than a purely postsynaptic formulation (37). We therefore propose that the classical scaling target is augmented by a term proportional to the AMPA-weighted afferent activity trace. This trace corresponds to the time-averaged AMPA-receptor-mediated current at the postsynaptic cell. Calcium influx through Ca^2+^-permeable AMPA receptors could make this signal accessible to the homeostatic machinery. The mismatch between the AMPA-weighted afferent activity trace and the postsynaptic activity trace is a candidate input to CaMKII. Frequency-dependent CaMKII activation could then drive the GluA1 trafficking that implements multiplicative afferent-weight rescaling. Once Hebbian STDP has built a heterogeneous per-synapse weight distribution, homeostasis tracks the AMPA-weighted afferent activity trace shaped by specifically potentiated inputs. The circuit can then relay firing rates selectively through learned pathways and restore that selectivity after perturbation. The biological timescale of these processes is hours to days. Our simulations converge over tens of stimulus epochs because molecular kinetics are abstracted into a per-epoch learning rate.

Three-factor learning rules such as e-prop and PP-prop depend on a spatially uniform per-layer eligibility trace amplitude for credit to flow coherently across depth (39, 40), a requirement that homeostatic scaling meets as a consequence of rate stabilization (Fig. 6). The same challenge, maintaining stable activation statistics across depth in the face of vanishing and exploding signals, motivated batch and layer normalization in artificial neural networks (22, 23). Our synaptic-scaling rule is a biologically plausible, local, and online counterpart to these global, batch-level techniques, operating per neuron with only locally available traces and without batching across examples. In this sense, relational homeostatic scaling plays the role of a local biological normalization rule, preserving activation scale across depth without batch statistics, global error signals, or externally tuned layer-wise gains. Two aspects of the biological solution may be informative for artificial normalization design. First, normalization is applied per neuron rather than per layer or per channel, allowing each unit to absorb its own intrinsic heterogeneity (Fig. 4). Second, the normalization target is a relationship between pre- and postsynaptic traces rather than a batch statistic of activations, so the rule does not require the population-level information that spiking neurons would not have access to. Whether per-neuron, trace-based normalization schemes can improve stability or sample efficiency in deep spiking networks remains an open question.

The model yields three experimentally testable predictions. First, pharmacological blockade of homeostatic scaling machinery, for example by disrupting TNF-*α* signaling or GluA1 trafficking, is predicted to impair inter-layer transmission of evoked firing rates even when within-layer rates are held in a normal range by E/I balance or other local mechanisms. Layer-to-layer rate consistency requires each feedforward link to operate near the critical per-synapse weight, and only the homeostatic total-weight constraint enforces this. Second, the equilibrium total afferent weight per neuron is predicted to correlate negatively with intrinsic excitability, with less excitable cells receiving stronger total synaptic drive at steady state, as predicted by 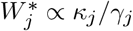 and illustrated in Fig. 4h. Third, after partial deafferentation or an abrupt reduction in afferent connectivity, the recovery timescale of inter-layer rate-code transmission is predicted to match the timescale of homeostatic scaling rather than that of Hebbian plasticity, because recovery acts through the total-weight constraint rather than the per-synapse weight structure.

Homeostatic synaptic scaling is typically described as a mechanism that restores circuit function after perturbation, but our results suggest that synaptic scaling may establish the operating regime in which rate-code transmission is possible in feedforward architectures. Associative plasticity then determines which signals are transmitted, and does so within the stable regime that relational homeostatic scaling autonomously maintains. Several limitations apply. The model is purely excitatory and feedforward. How the relational rule interacts with inhibitory plasticity in recurrent networks, and whether an analogous rule on inhibitory synapses could jointly stabilize E/I-balanced recurrent circuits, remain open questions. The training protocol uses a fixed stimulus rate per epoch, and whether the total-weight constraint remains effective under naturalistic, time-varying afferent patterns remains to be shown. The learning rate *η* is accelerated relative to the biological timescale of synaptic scaling, and the qualitative conclusions depend only on the separation between scaling and Hebbian timescales, not on their absolute values.

## Materials and Methods

### Network and neuron model

We consider a purely feedforward stack of *L* = 10 layers of current-based leaky integrate-and-fire (LIF) units, driven by an input layer of *N*_in_ = 1000 independent Poisson spike generators firing at rate *r*_in_. Each hidden layer contains *N* = 1000 neurons and receives afferents from the preceding layer through a fixed Bernoulli mask 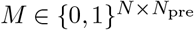 of density *p*, so that the expected fan-in is *K* = *N*_pre_ *p*. Membrane and synaptic dynamics for neuron *j* obey

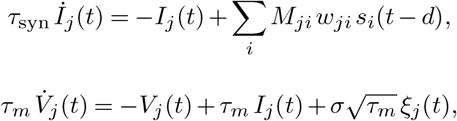

where 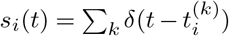 is the afferent spike train, *d* a conduction delay, and *ξ*_*j*_ independent white noise. A spike is emitted whenever *V*_*j*_(*t*^−^) ≥ *V*_th_, after which *V*_*j*_ is clamped to the reset *V*_reset_ = 0 for a refractory interval *τ*_ref_. All voltages are dimensionless (*V*_th_ = 1). Unless otherwise stated, *τ*_*m*_ = 20 ms, *τ*_syn_ = 10 ms, *τ*_ref_ = 5 ms, *d* = 2 ms, *σ* = 0.15, *p* = 0.2, and *r*_in_ = 30 Hz.

### Heterogeneity

Each neuron carries a gain factor *κ*_*j*_ and a membrane time constant *τ*_*m,j*_, sampled i.i.d. from a distribution with mean equal to the homogeneous reference and coefficient of variation *σ*_*κ*_ or 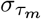. Four distribution families are considered (normal truncated at zero, uniform, lognormal, and gamma), each reparameterized to share the prescribed mean and coefficient of variation.

### Plasticity rules

Let *W*_*j*_ = ∑_*i*_ *w*_*ji*_ denote the total excitatory weight onto neuron *j*. Plasticity is defined on three families of local rules.

#### Relational homeostatic scaling

Each neuron tracks two low-pass traces with common time constant *τ*_*r*_,

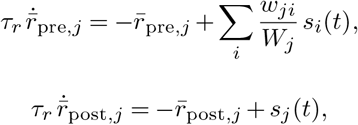

so 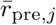 is the AMPA-weighted afferent activity trace and 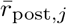 is the postsynaptic activity trace. The total afferent weight evolves as

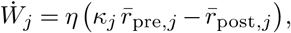

and is implemented by multiplicative rescaling of all excitatory afferents onto neuron *j*, so their relative strengths are preserved (34). The fixed point is 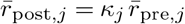, meaning that each neuron aligns its postsynaptic activity trace with its AMPA-weighted afferent activity trace.

#### Target-rate fixpoint

An alternative homeostatic term drives 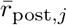 toward a fixed target *r*_tgt_ through 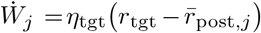 . This term can be added to the relational rule to yield the combined rule used in Fig. 2.

#### Total-weight normalization

The total excitatory afferent weight *W*_*j*_ onto each neuron is continuously held at its initial value, locking the total afferent weight while leaving STDP free to redistribute individual synaptic strengths (33, 34).

#### Spike-timing-dependent plasticity

STDP adjusts synaptic strengths based on the relative timing of pre- and postsynaptic spikes, with causal pairings (postsynaptic after presynaptic) driving potentiation and anti-causal pairings driving depression in the pair-based variant (46). Each synapse carries a fast pre-synaptic eligibility trace 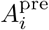 and a fast post-synaptic trace 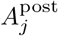, both decaying with time constant *τ*_±_,

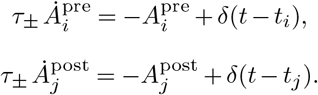

On a presynaptic spike at time *t*_*i*_, the weight receives a depression increment 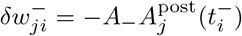. On a post-synaptic spike at time *t*_*j*_, the weight receives a potentiation increment 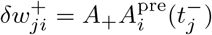, with *A*_+_ = 2 × 10^−4^ and *A*_−_ = *A*_+_ · (*τ*_+_/*τ*_−_) · 1.10 ≈ 2.2 × 10^−4^. Anti-causal STDP reverses both signs. Symmetric STDP sets *A*_−_ < 0 so that both coincidence directions are potentiating. The triplet rule of Pfister & Gerstner (47) supplements the pair kernel with a slow pre-synaptic trace 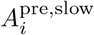 (decay 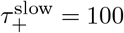 ms) and a slow post-synaptic trace 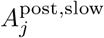 (decay 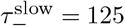 ms). The depression amplitude on a presynaptic spike becomes 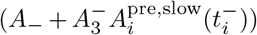, and the potentiation amplitude on a postsynaptic spike becomes 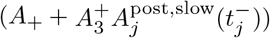, with 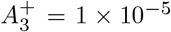 and 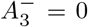. All increments are accumulated over the stimulus epoch, summed, and applied once at epoch end. Weights are clipped to *w*_*ji*_ ≥ 0 after the update. When combined with relational homeostatic scaling, STDP drives per-synapse weight structure while the multiplicative afferent rescaling constrains *W*_*j*_.

### Simulation and training protocol

All simulations are implemented in PyTorch with 1 ms discrete time steps and no automatic differentiation. LIF dynamics, eligibility traces, and plasticity rules are integrated by explicit Euler updates with exponential-decay coefficients.

Each training epoch presents a 500 ms stimulus in which *N*_in_ = 1000 independent Poisson spike generators fire at *r*_in_ = 30 Hz (1). During the epoch, the relational traces 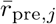 and 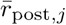 are updated online at every millisecond, and STDP eligibility traces accumulate spike-pair increments continuously. At epoch end, the STDP update is applied first (accumulated increments are added to each weight, then clipped to zero), followed by the homeostatic multiplicative afferent rescaling, so STDP reshapes relative synaptic strengths before homeostasis renormalizes the total afferent weight.

The homeostasis learning rate is *η* = 0.1 and the target-rate term learning rate is *η*_tgt_ = 10^−4^. The relational trace time constant is *τ*_*r*_ = 200 ms. Fast STDP traces decay with *τ*_±_ = 20 ms. The triplet rule additionally uses slow traces with 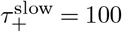 ms and 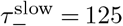 ms. STDP amplitudes are 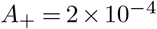 (potentiation) and *A*_−_ ≈ 2.2 × 10^−4^ (depression). The triplet rule additionally uses 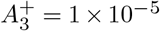 and 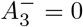. Initial per-synapse weights are uniform over connected pairs at *w*_0_, giving an initial total afferent weight 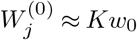.

For the single-layer frequency-generalization analysis, a layer was trained for 100 epochs at 30 Hz from *w*_0_ = 0.005. The trained weights were frozen. The input rate was then swept from 5 to 80 Hz in 5-Hz steps with three random seeds per rate.

### Mean-field rate theory and critical weight

In a mean-field approximation, each neuron is described by the transfer function *r* = *ϕ*(*Wr*_in_), and the gain of layer *ℓ* for small rate perturbations around an operating point is 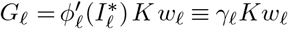. Stable rate-code transmission across depth requires *G*_*ℓ*_ = 1, which defines a critical per-synapse weight *w*^∗^ 1*/*(*γ*_*ℓ*_*K*). Predictions in Figs. 1 and 3 were obtained by numerical mean-field simulation of the single-neuron LIF dynamics under Poisson input and bracketing the *w*-value at which unit gain is achieved.

Linearizing the relational homeostatic dynamics around this operating point yields a linear ODE in *W*_*j*_ with fixed point 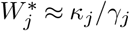 and relaxation time 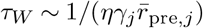, so convergence is faster at higher presynaptic activity or neuronal gain.

### Eligibility traces for local credit assignment

As a diagnostic of downstream learnability, each layer maintains two low-pass traces with time constant *τ*_pp_ = 50 ms (39, 40),

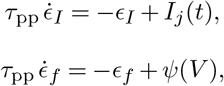

where *ψ*(*V*) = *γ*_surr_ max(0, 1−|*V*−*V*_th_|*/V*_th_) with *γ*_surr_ = 0.3 is the piecewise-linear surrogate gradient (48) used in spiking-network learning. The per-layer eligibility amplitude reported in Fig. 6 is ∥*ϵ*_*f*_∥ ∥*ϵ*_*I*_∥ averaged over the stimulus window. These traces are read out diagnostically and do not enter the plasticity update.

## Author contributions

Huihong Li: Conceptualization, Methodology, Software, Formal Analysis, Writing – Original Draft, Visualization. Robert A. McDougal: Conceptualization, Supervision, Writing – Review & Editing, Funding Acquisition.

## Acknowledgments

We thank our lab members for engaging discussions and improvements of the manuscript.

## Supplementary Figure

**Fig S1.**
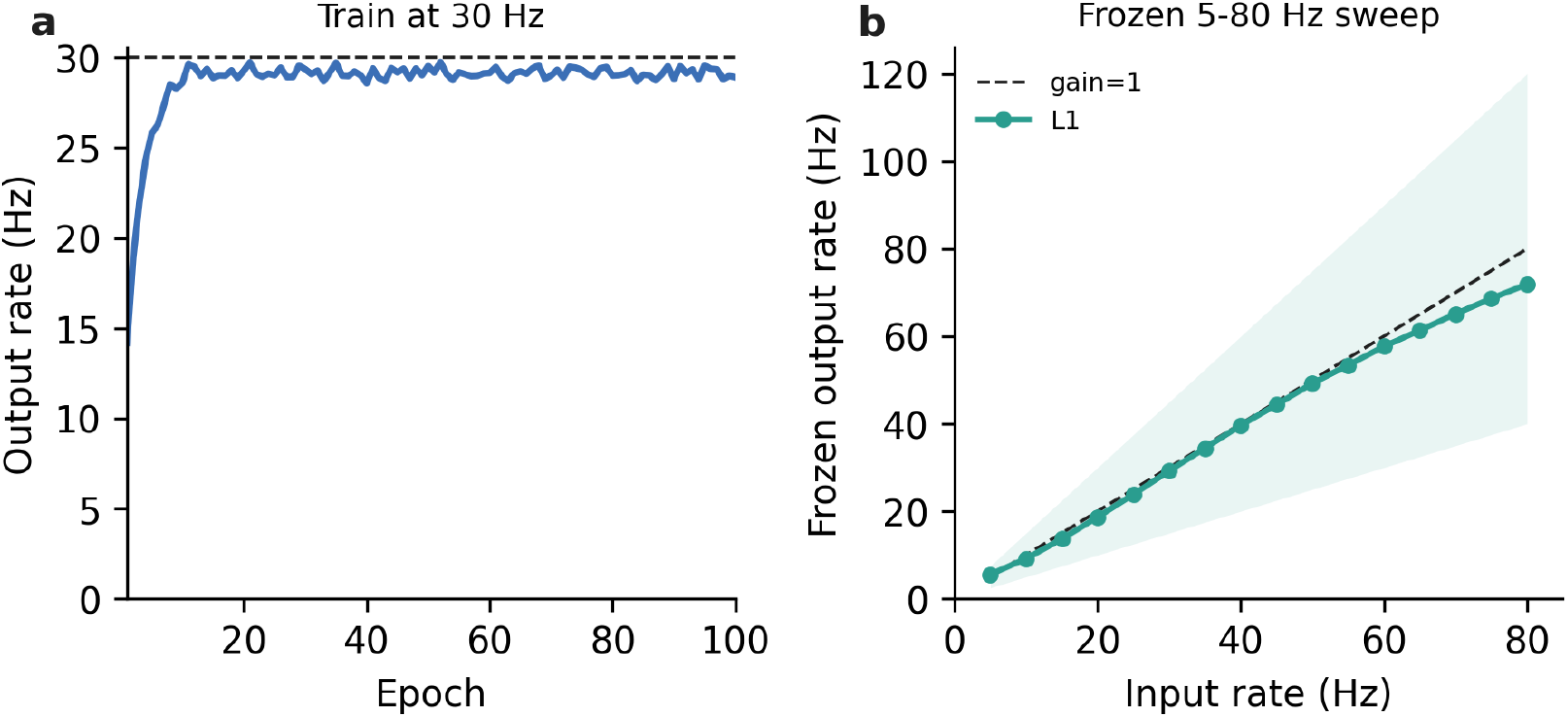
A layer trained at 30 Hz generalizes across input rates. A single LIF layer was trained for 100 epochs from the decay initialization with relational homeostatic scaling under a 30-Hz Poisson input. The trained weights were frozen, and the Poisson input rate was swept from 5 to 80 Hz. (a) The training trajectory converged to the 30-Hz target. (b) The frozen output rate tracked the input rate across the sweep. The shaded band marks gain between 0.5 and 1.5.

